# Dendritic excitability controls overdispersion

**DOI:** 10.1101/2022.11.18.517108

**Authors:** Zachary Friedenberger, Richard Naud

**Affiliations:** Centre for Neural Dynamics and Artificial Intelligence, University of Ottawa, Ottawa ON, Canada; Department of Physics, University of Ottawa, Ottawa ON, Canada; Department of Cellular and Molecular Medicine, University of Ottawa, Ottawa ON, Canada

## Abstract

The brain is an intricate assembly of intercommunicating neurons whose input-output function is only partially understood. The role of active dendrites in shaping spiking responses, in particular, is unclear. Although existing models account for active dendrites and spiking responses, they are too complex to analyze analytically and demand long stochastic simulations. Here we combined cable and renewal theory to describe how input fluctuations shape the response of neuronal ensembles with active dendrites. We found that dendritic input readily and potently controls interspike interval dispersion. This phenomenon can be understood by considering that neurons display three fundamental operating regimes: one mean-driven regime and two fluctuation-driven regimes. We show that these results are expected to appear for a wide range of dendritic properties and verify the predictions of the model in experimental data. These findings have implications for the role of interspike interval dispersion in learning and for theories of attractor states.

## INTRODUCTION

Understanding computation within neuronal networks relies crucially on grasping how the inputs to a neuron affect its output [1–6]. The canonical input-output function is assumed to produce i) firing-frequency increasing monotonically with input intensity and ii) a concomitant decrease of interspike interval dispersion as measured through the coefficient of variation (CV) [7–9]. These properties arise from a transition between two fundamentally different regimes: the fluctuation-driven regime where CV is high but firing frequency is modest and the mean-driven regime where firing frequency is high but the CV is low [7–9]. Yet, this view of the input-output function of neurons remains an assumption, one that is based on the predominance of evidence about the integrative properties of the cell body [10–12].

Nonlinear properties of dendrites are expected to profoundly influence the input-output function [13, 14]. Calcium, NMDA and sodium spikes are widespread instances of such nonlinear properties [15–21]. Researchers have hypothesized that they modulate the gain of the soma-based input-output function [22, 23] or implement a cascade of nonlinear transformations with additive interactions across units, as in an artificial neural network [24– 30]. However, the effects that elicited these hypotheses appear weaker in the presence of background fluctuations [22, 26]. Alternatively, dendrites may alter the input-output function less in terms of the firing frequency (the f-I curve), and more in terms of the patterns of interspike intervals (the CV-I curve). Consistently, dendritic spikes can control the generation of bursts of action potentials [16, 31, 32], shaping interval dispersion.

To study the integrative properties of dendrites and the soma in the presence of noise, we integrated elements of cable theory [33, 34] into a leaky integrate-and-fire (LIF) framework [35–37] permissive to the direct computation of response statistics (renewal theory [5, 38–40]). This technique permits computation of the interspike interval distribution of an ensemble of neurons with active dendrites, without relying on subdivision of compartments [41] nor stochastic sampling [23]. We found that, contrary to dendrite-less models, input intensity to both the dendrites and the soma can increase the CV.

## RESULTS

We consider a ball-and-stick model consisting of a cell body connected to a uniform yet excitable dendrite. Dendritic spikes (*η*_*d*_(*x, t*)) can affect the somatic potential and a spike in the soma can affect both the refractory effects in the soma and the backpropagating action potential in the dendrite (*η*_*s*_(*x, t*); Fig. 1a). The spatiotemporal membrane potential at the soma (at *x* = 0) and across the dendrite (*x >* 0) resulting from the spatiotemporal input *I*(*x, t*) can be written as,

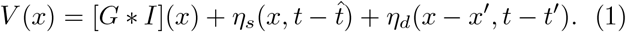

Where *G*(*x*) is the Green’s function of the passive cable [33, 34], 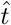 is the time of the last somatic spike, *x*^*′*^, *t*^*′*^ is the time and point of origin of the last dendritic spike. Combining this spatio-temporal neuron model with stochastic firing conditions and renewal theory allows us to compute the firing rate and CV for stationary inputs in the presence of noise (see Methods).

**FIG. 1.**
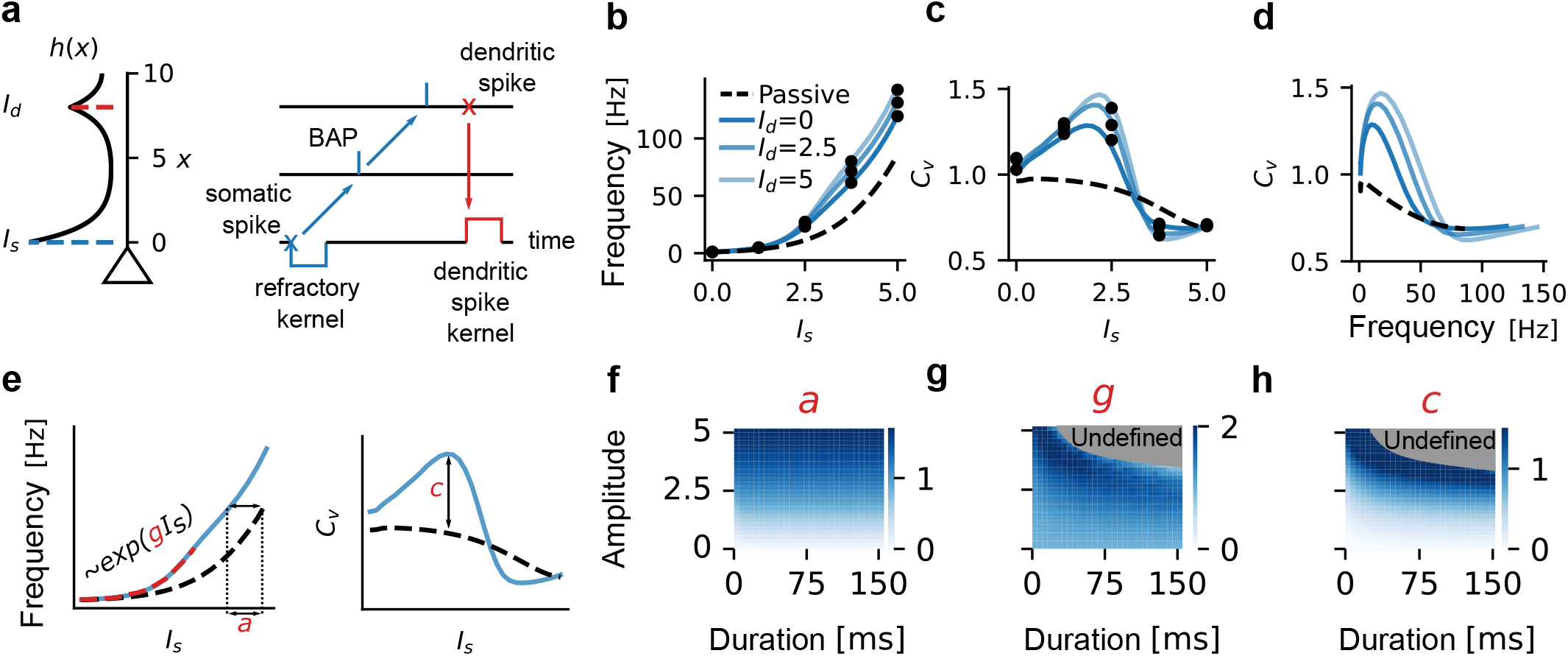
Dendritic input controls overdispersion. **a** Left: Stationary input potential resulting from the Green’s function of a ball-and-stick model receiving localized input currents at the soma (*x* = 0, *I*_*s*_ = 5) and in the dendrite (*x* = 8, *I*_*d*_ = 5). Right: Schematic illustration of spike-triggered effects. A somatic spike causes a refractory period in the soma and a backpropagating action potential (BAP) along the dendrite. A dendritic spike creates a short-lived depolarization in the soma. **b** Firing rate and **c** coefficient of variation against somatic input current for three dendritic input strengths. The black dashed line corresponds to the absence of dendritic spikes. The black dots correspond to points obtained using Monte Carlo simulation (see Methods). **d** CV against firing frequency for three dendritic input strengths. **e** Schematic of the parametrization of dendritic input modulation in terms of the shift (*a*) and gain (*g*) of the f-I curve, as well as overdispersion (*c*). **f-h** Heatmaps of the effect on shift (f), gain (g) and overdispersion (h) for different amplitude and duration of the dendritic spikes. The undefined region in (g) and (c), corresponds to a region where our definitions of gain and dispersion break down (see Methods).

For simplicity, we focused on a single dendritic input that is electronically distant from the cell body so that it does not affect it directly. The Green’s function spreads the effect of this input along the dendrite (Fig. 1a). We build f-I and CV-I curves by varying the intensity of the somatic input for different intensity of the dendritic input (Fig. 1b-d). As expected, f-I curves were modulated by the strength of dendritic inputs in two fundamental ways. Firstly, the f-I curves shift by an amount that scales with dendritic input strength, consistent with dendrites acting additively such as in hierarchical neural networks [14, 26, 28]. Secondly, the gain of the f-I curve increased slightly with dendritic input, consistent with a role of dendritic spikes in gain modulation [22, 23]. Averaging 10 000 Monte-Carlo simulations per point corroborates our formalism (black dots in Fig. 1a) as well as the effects on f-I curve gain and offset.

In the LIF receiving Gaussian-distributed inputs, CVs are smaller or equal to one and monotonically decrease with the strength of somatic input (Fig. 1c-d) [7, 8]. In the presence of dendritic excitability, however, the CV shows overdispersion in a manner controlled by the strength of dendritic and somatic inputs. The CV-I relationship becomes non-monotonic: increases in somatic input strength increase dispersion before decreasing it. Strikingly, the most salient part of the CV modulation occurs for more moderate somatic inputs than both gain and additive modulation, in a firing regime consistent with in vivo condition. We also extended the theory to take into account hotspots for dendritic spike generation and show CV modulation arises whenever dendritic inputs target the hotspot (Fig. S1).

To establish the generality of this phenomenon, we exploited the computational efficiency of our approach to perform parameter sweeps. We focused on the amplitude and duration of dendritic spikes as seen in the soma (Fig. 1e-h). For each set of parameters, we computed the additive modulation, the gain modulation, and the CV modulation (see Methods). We found that CV modulation occurs on a wide range of parameters, and is stronger for moderately large dendritic spike amplitudes and durations, an expression pattern echoed by the effects on gain, albeit with diminished potency. Divergently, additive modulation is solely controlled by dendritic spike amplitude. Overall, we find that CV modulation by dendritic inputs occurs for a large range of parameters.

How can we understand the non-monotonic relationship between CV and I? In point neuron models, the transition between high and low dispersion is understood in terms of the transition between fluctuation- and mean-driven regimes, defined as whether the mean input reaches threshold or not. Dendritic spikes, perceived as sustained depolarizations, can similarly either reach the threshold or not, depending on the mean somatic input and the amplitude of dendritic spikes. To illustrate this in a simple model, we simulate a two-compartment model with a LIF soma (Fig. 2a-b). When the mean somatic input was low such that dendritic spikes remain subthreshold, an increase of the dispersion was achieved by switching between two states having high variability of interspike intervals (a Cox process; Fig. 2c). When the somatic input was sufficiently strong, dendritic spikes will on average cross the threshold leading to more regular intervals in one of the two states (inducing burst-like events; Fig. 2d). As the mean somatic input increases, it forces the dendritic spikes to cross threshold and thus switches the Cox-process to a burst-like process. Higher somatic input continues to decrease the CV by entering the mean-driven regime (Fig. 2e). According to this theory, the transition between the Cox-process regime and the burst-like regime depends on the amplitude of the dendritic spike as seen from the soma. We verified that increasing the amplitude of dendritic spikes decreases the somatic current for which CV is maximum (Fig. 2f). Thus, the input-output function of a neuron can be separated in three fundamentally distinct regimes with distinct dependence on somatic and dendritic inputs (Fig. 2c): i) a Cox-process regime where neither dendritic spikes nor the mean input reach threshold in the absence of fluctuations, ii) a burst-like regime where dendritic spikes reach threshold in the absence of fluctuations but the mean somatic input does not, and iii) a mean-driven regime where the mean is above threshold.

**FIG. 2.**
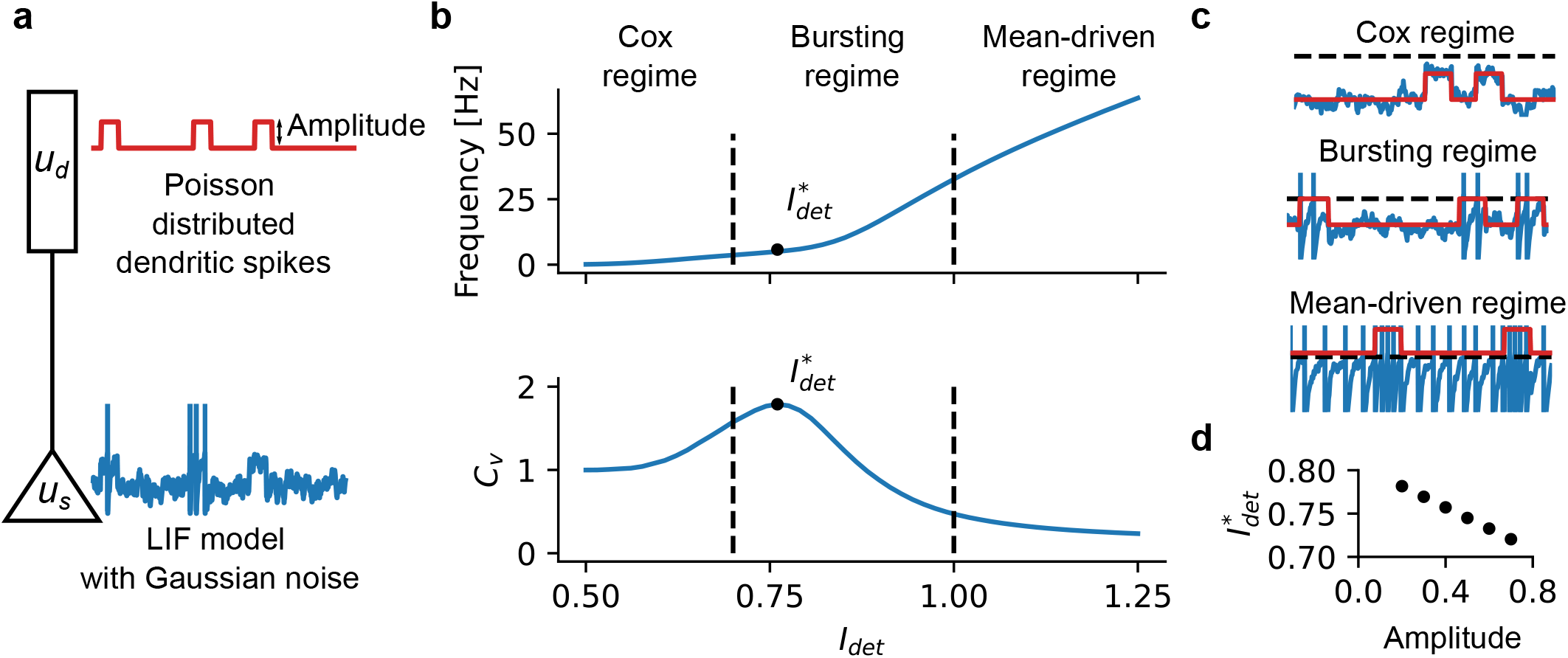
Three operational regimes. **a** Schematic of two-compartment model with noisy input injections. **b** f-I and CV-I curves highlighting the transition between three operational regimes: Cox regime where mean input plus dendritic spike amplitude is below threshold, Bursting regime where mean input plus dendritic spike amplitude is above threshold, Mean-driven where average input alone is above threshold. **c** Voltage traces corresponding to the three regimes. **d** The strength of the somatic input associated with the peak CV 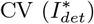 shown against amplitude of the dendritic spike.

There arise one core prediction: CV of interspike intervals should increase when mean input to dendrites is increased. This effect is likely to be observed more strongly at lower firing rates. To test this prediction, we used previously published dual patch clamp experiments with noisy current injection simultaneously in the soma and the apical dendrite [22]. Fig. 3 shows both the f-I curves and the CV-I curves for increasing mean dendritic inputs. While a combination of gain and additive modulation is visible from the f-I curves, the CV-I curves show that increasing dendritic inputs increases the CV at low firing rates, verifying this core prediction. Another prediction is that for a neuron in vivo, synaptic input to the dendrite should reach higher dispersion than current injection to the cell body. This has been observed in visual cortex [42] where current stimulation in vivo has been compared with sensory stimulation. Together, the dendritic control of overdispersion is an experimentally observed phenomenon that is likely to take place whenever sustained and potent dendritic spikes affect the cell body.

**FIG. 3.**
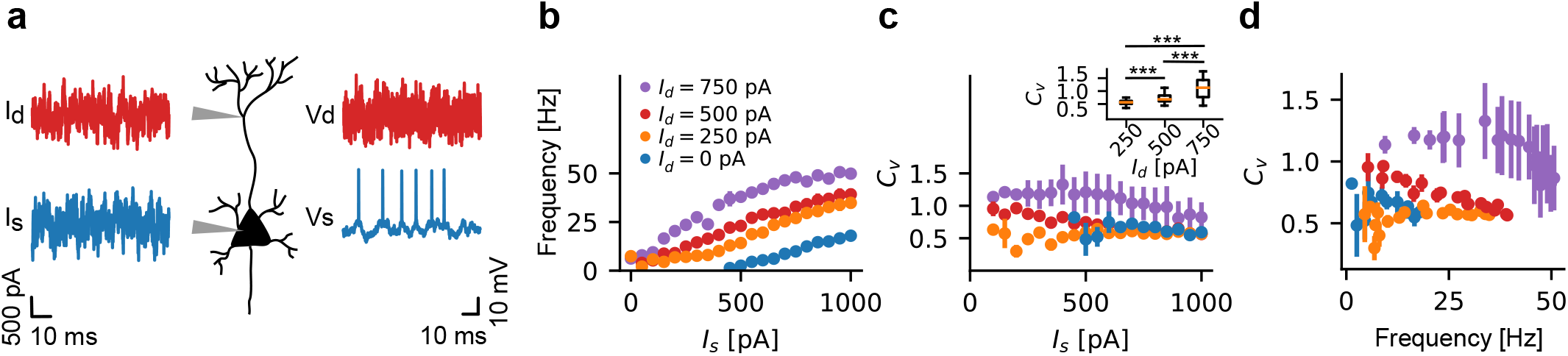
Experimental validation. **a** Schematic of experiment [22]: dual patch clamp was performed and noisy currents were injected in the soma and the apical dendrite. **b** Time-averaged firing rate is shown against the mean strength of the somatic input for four levels (0, 250, 500, 750 pA) of dendritic input (blue, orange, red, purple, respectively). **c** CV as a function of somatic and dendritic input strength (same color code as a). CV pooled over all somatic input conditions and analyzed with Welch’s unequal variances t-test (p*<*0.001, after correcting for multiple comparisons). **d** CV against mean activity for the four levels of dendritic input. Error bars represent the standard error in the mean.

## DISCUSSION

We have introduced a computational technique able to directly compute statistics from an ensemble of neurons with active dendrites. This theory can be generalized to study time-dependent inputs [39, 40], network interactions [5] and pairwise correlations [43].

We found that in the burst-like regime, interval dispersion is mostly controlled by the strength of dendritic inputs. Multiplexing, whereby somatic and dendritic inputs are both decodable from neuronal responses [31, 44], is thus expected to be most reliable in this regime. This suggests that the principle of multiplexing as a solution to the credit-assignment problem [45] is likely restricted to cell subtypes having dendritic spikes of large amplitude.

Overdispersion is widely observed in cortex [42, 46], even in conditions of sustained firing associated with working memory [47]. Although existing models could generate high dispersion, they did so at the cost of either excessively large input noise [9] or required a fine-tuned attractor state [3, 48]. Our results suggest that dendritic spikes are a potential solution to this long-standing problem.

## METHODS

### A stochastic spatiotemporal model with active dendrites

We consider a spatiotemporal membrane potential, *V* (*x, t*), arising from a combination of three different processes: 1) the passive integration of inputs *h*(*x*), 2) spatiotemporal effects of a somatic action potential at time 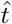, and 3) of a dendritic action potential at time *t*^*′*^ and location *x*^*′*^. These three processes are added linearly to give,

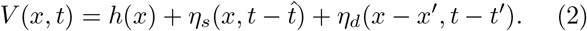

In general, the input potential along a passive dendrite *h*(*x, t*) is given by the cable equation,

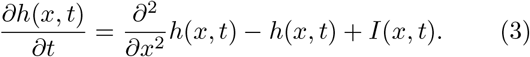

Where *x* and *t* are dimensionless variables for space and time respectively. The scaling from variables 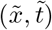 with proper dimensions is given by, 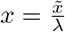, and 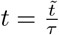. Where 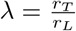 is the characteristic electronic length scale given by the ratio between the transversal current per unit length *r*_*T*_ and the longitudinal current per unit length *r*_*L*_. The characteristic time scale, *τ*, is given by the product of *r*_*T*_ and the capacitance per unit length *c, τ* = *r*_*T*_ *c*. In eq. (3), the external input current *I*(*x, t*) has been scaled to have the same units as *h*(*x, t*), with the scaling from proper units *Ĩ* given by *I* = *r*_*T*_ *Ĩ*.

For a spatially detailed but temporally stationary input *I*(*x*), the solution to equation (3) is given by the convolution,

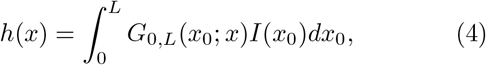

where *G*_0,*L*_(*x*_0_; *x*) is called the Green’s function. For a passive cable of length *L* it can be found explicitly [34],

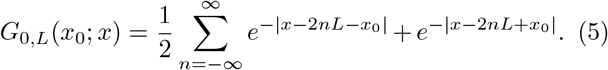

Therefore, given an arbitrary input current *I*(*x*), we can determine the input potential *h*(*x*) for equation (2) using equations (4) and (5). In Fig. 1a, *h*(*x*) is shown for an input of *I*(*x*) = *I*_*s*_*δ*(*x*) + *I*_*d*_*δ*(*x* − 8), where *δ*(*x*) is the Dirac delta function.

The kernel *η*_*s*_(*x, t*) in equation (2) captures the stereotypical effects of refractoriness and adaptation [35, 37] at the soma, and the effect of the backpropagating action potential in the dendrites [16, 49, 50]. We let the somatic kernel correspond to a simple rectangular refractory period combined with a back-propagating action potential modelled as a traveling depolarizing pulse. Specifically, a somatic spike at time 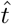 changes the membrane potential by,

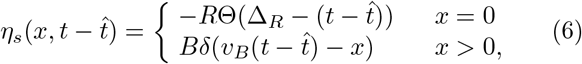

where Θ(*x*) is the Heaviside step function, *R* is the amplitude of the refractory period, Δ_*R*_ is the refractory period duration, *B* is the amplitude of the backpropagating action potential, and *v*_*B*_ is the speed of the backpropagating action potential.

Similarly, the kernel *η*_*d*_(*x, t*) in equation (2) captures the stereotypical effects of having emitted a dendritic spike. We note that most of the dendritic spikes do not have a stereotypical spatio-temporal effect on the neuron, since both calcium and NMDA spikes display variable durations [15, 19, 51]. This formulation is, therefore, an approximation corresponding to an average time-course and spatial extent. To simplify the dynamics of the neuron model, we assume that influence of the dendritic spike at the soma is instantaneous and independent of the den-dritic spike location *x*^*′*^ and we neglect the local effects of the dendritic spike. Therefore, we model the influence of the dendritic spike at the soma as a rectangular pulse with amplitude *D* and duration Δ_*D*_. A dendritic spike at location *x*^*′*^ and time *t*^*′*^ thus changes the membrane potential by,

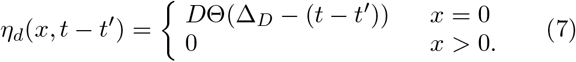

To be complete, the neuron model requires conditions for the emission of somatic and dendritic action potentials. Although these can be deterministic, we seek a stochastic description in order to take into account the variability of ion channels, synaptic transmission and network activity. As in previous descriptions [12, 35, 37], we take the instantaneous hazard rate *λ*_*s*_ for emitting a somatic spike to be an exponential readout of the membrane potential at the soma,

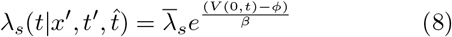

where 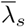 is a parameter that defines the baseline firing rate at threshold *ϕ*, and where *β* scales the noise of the stochastic emission. For dendritic spike emission, we take the hazard rate at location *x* and time *t* to be a rectified linear function of the membrane potential along the dendrite,

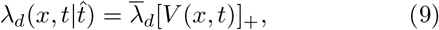

where [*V*]_+_ = 0 if *V*) ≤ 0 and [*V*]_+_ = *V* if *V >* 0.

To simplify the survivor function of our neuron model, which will be derived in the next section, we assume that the soma forgets the effect of the last dendritic spike after each new somatic spike. That is, the somatic hazard function only depends on the last somatic spike 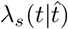 if 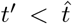, but also depends on the last dendritic spike 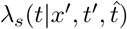 if 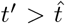. This can be thought of as the effect of the dendritic spike disappearing immediately after each somatic spike because of the reset of the membrane potential. For clarity, from this point forward we differentiate between these hazards by defining the initial so-matic hazard rate 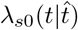 for 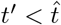, and the conditional somatic hazard rate 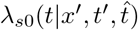 for 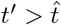.

Using equations (4), (6), and (8), we define the initial somatic hazard rate to be,

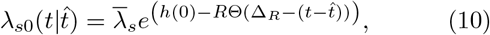

where we have absorbed the threshold *ϕ* into the constant 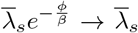 and the parameters *I*_*s*_, *I*_*d*_, *R*, and *D* have been scaled by 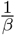, making them dimensionless variables. Similarly, using equations (4), (6), (7), and (8), we define the conditional somatic hazard rate to be,

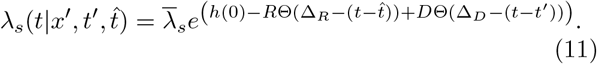

Lastly, using equations (4), (6), and (9), we define the dendritic hazard rate to be,

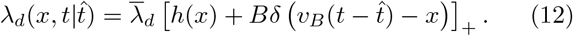

### Dendritic renewal theory

To derive the stationary firing rate *A*_*∞*_ (which we refer to as the frequency in the figures and main text) and coefficient of variation *C*_*v*_ for an ensemble of neurons with active dendrites, we use the firing conditions of the neuron model defined by equations (10), (11), and (12) and adapt renewal theory [5, 38–40] to a spatio-temporal event generation model.

Given the survivor function of a renewal process, 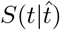, which gives the probability that the neuron model hasn’t fired a spike up and until time *t* given the last spike occurred at time 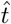, we can calculate the stationary firing rate,

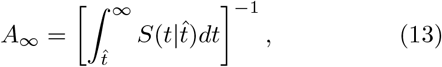

and the coefficient of variation of the inter-spike-interval (ISI) distribution,

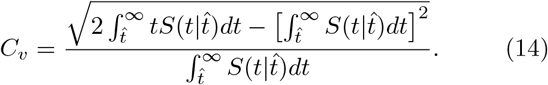

Therefore, our goal is to derive an approximation of 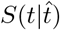 for our neuron model, which we call the marginal somatic survivor function, as the dependence on dendritic firing will be marginalized out.

To determine 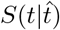, we use the hazard functions from equations (10), (11), and (12) to define three survivor functions. The dendritic survivor function 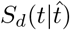, the initial somatic survivor function 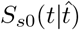, and the conditional somatic survivor function 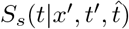. Which are given by,

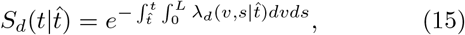

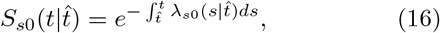

and

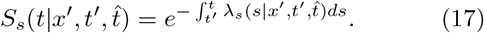

Given the dendrite is free to spike an arbitrary number of times during each somatic ISI and the somatic hazard function is momentarily increased following each dendritic spike, the marginal somatic survivor probability will depend on the probability of all possible future dendritic spikes occurring during the somatic ISI. Therefore, the marginal somatic survivor function can be obtained by path integration. That is, adding up all the possible paths through which the soma survives from 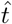 up to time *t* without firing a spike. However, the full path integral is intractable for even simple choices of dendritic and somatic hazard rates, so we consider the case where there is at most one dendritic spike during each somatic ISI. This can be thought of as an approximation where the dendritic firing rate is less than or equal to the somatic firing rate. With this additional assumption, the marginal survivor probability can be written as,

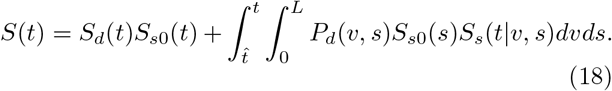

Where we have dropped the dependence on 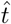 to simplify notation and we have introduced *P*_*d*_(*x*^*′*^, *t*^*′*^), which is the spatio-temporal dendritic interval distribution, given by *P*_*d*_(*x*^*′*^, *t*^*′*^) = *λ*_*d*_(*x*^*′*^, *t*^*′*^)*S*_*d*_(*t*^*′*^). The first term *S*_*d*_(*t*)*S*_*s*0_(*t*) can be interpreted as the probability that both the soma and the dendrite independently surviving until time *t* given the last somatic spike was at 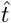. The second term *P*_*d*_(*x*^*′*^, *t*^*′*^)*S*_*s*0_(*t*^*′*^)*S*_*s*_(*t*|*x*^*′*^, *t*^*′*^)*dx*^*′*^*dt*^*′*^ is the probability that the soma and the dendrite both survive up until time *t*^*′*^, the dendrite fires a spike at (*x*^*′*^, *t*^*′*^), and then the soma survives until time *t* given the dendritic spike occurred at (*x*^*′*^, *t*^*′*^). The integral then runs over all possible dendritic spike locations *x*^*′*^ and times *t*^*′*^. To include the possibility of more dendritic spike occurring during the somatic ISI, additional terms can be added to equation (18), integrating over all combinations of dendritic spike times.

### Monte Carlo simulation

To verify our integration approach we used Monte Carlo simulations with the same restrictions on spike generation. At each time step, we calculate the probability of firing a somatic spike in the interval [*t, t* + Δ*t*] using the somatic hazard rates,

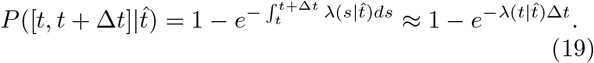

If no dendritic spike has occurred since the last somatic spike, 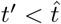, then the initial somatic hazard rate 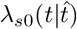 is used. If a dendritic spike has occurred since the last somatic spike, 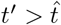, then the conditional somatic hazard rate 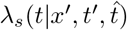 is used instead.

To sample the dendritc spike times and their location along the dendrite, we write the joint location-interval distribution using the product rule, 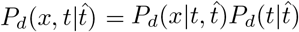. The marginal distribution 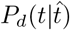 is given by, 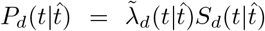. Where 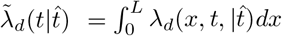 and 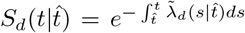. The condi-tional density for dendritic spike location is then given by,

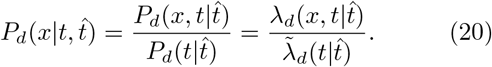

We can then simulate the process by iterating through the discretized values for *t* = {Δ*t*, 2Δ*t*, …, *N* Δ*t*}. The probability of emitting a dendritic spike in the interval [*t, t* + Δ*t*] is then given by equation (19) with 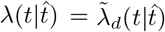. If a spike occurs, we then sample from the condi-tional distribution 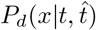. However, because our dendritic hazard rate from Eq. (12) has a delta function located at *x* = *v*_*b*_*t*, our conditional density is a mixture of a continuous probability density and a discrete probability. To properly sample this mixture of distributions, we note that the integral over *x* is normalized, such that 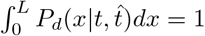, which gives us,

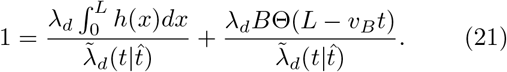

The first term on the right is the total probability of the continuous part and the second term is the discrete probability of observing a dendritic spike at point (*x, t*) = (*v*_*B*_*t, t*). If a dendritic spike occurs at time *t*^*′*^, we first sample a uniform random number *r*_1_ ∼ *U* (0, 1). If 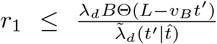, then the location of the den-dritic spike is *x*^*′*^ = *v*_*B*_*t*^*′*^, occurring at the location of the back propagating action potential. However, if 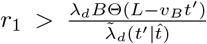, we then use rejection sampling on the continuous part of the distribution 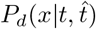. That is, we sample a prospective location at random from a spatial grid, *x*^*′*^ ∼ {0, Δ*x*, 2Δ*x*, …, *L* − Δ*x*} and another uniform random number *r*_2_ ∼ *U* (0, 1). If 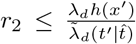 we accept *x*^*′*^ as the location of the dendritic spike. Oth-erwise, *x*^*′*^ is rejected and we resample *x*^*′*^ and *r*_2_ until a location is accepted.

For each input value pair (*I*_*s*_, *I*_*d*_), we ran the Monte Carlo simulation 10, 000 times. Each run had a total length of 3, 000 steps with a step size Δ*t* = 0.01, which corresponds to 1 ms steps because our time constant was chosen to be *τ*_*m*_ = 10 ms. The spatial grid spacing was defined to be, Δ*x* = *v*_*B*_Δ*t*, where *v*_*B*_ is the speed of the backpropagating action potential.

### Leaky integrate-and-fire neuron with dendritic spikes

We model a leaky integrate-and-fire neuron with white noise somatic input and dendritic spikes acting as stochastic increases in membrane potential. The somatic membrane potential is given by the stochastic differential equation,

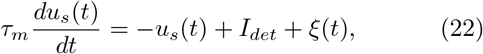

where *ξ*(*t*) is a white noise input with ⟨*ξ*(*t*)⟩ = 0 and ⟨*ξ*(*t*)*ξ*(*t*^*′*^)⟩ = *σ*^2^*τ*_*m*_*δ*(*t* − *t*^*′*^). *I*_*det*_ is a constant deterministic input. Dendritic spikes occurring at times 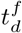 were distributed according to a Poisson process with rate *λ*_*d*_. Each dendritic spike then caused a temporary increase in the membrane potential, 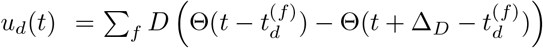. Where *D* is the amplitude and Δ_*D*_ is the duration of the dendritic spike. Somatic spike times *t*_*s*_ were recorded whenever *u*_*s*_(*t*_*s*_) + *u*_*d*_(*t*_*s*_) ≥ 1. Then *u*_*s*_(*t*) was reset to zero the following time step, *u*_*s*_(*t*_*s*_ + *dt*) = 0.

### Numerical methods

In Fig. 1b-d, the input is given by *I*(*x*) = *I*_*s*_*δ*(*x*) + *I*_*d*_*δ*(*x* − 8) along a dendrite of length *L* = 10. The somatic input *I*_*s*_ and dendritic input *I*_*d*_ levels were varied and numerical integration was used to find *A*_*∞*_ and *C*_*v*_ using equations (13) and (14). The membrane time constant was chosen to be *τ*_*m*_ = 10 ms. The other neuron model parameters were fixed to be, {*R* = 1, Δ_*R*_ = 10 ms, Δ_*D*_ = 20 ms, *v*_*B*_ = 2, *B* = 1, *λ*_*s*_ = 0.01, *λ*_*d*_ = 0.1}. For the case of an active dendrite, the dendritic spike amplitude was set to be *D* = 2, while the amplitude was set to *D* = 0 for the case of a passive dendrite.

In Fig. 1f-h, simulations to obtain the f-I and CV-I curves were repeated with the same parameters, expect the dendritic spike amplitude *D* and duration Δ_*d*_ were also varied and the dendritic current was fixed to be *I*_*d*_ = 2.5. The additive component, *a*, as shown in Fig 1e and Fig 1f, was defined as the absolute change in *I*_*s*_, between the active and passive condition, required for the f-I curve to reach 80 Hz. As a result of the dendritic spike, the log transformed f-I curve in the active condition could be captured by a piece-wise linear function given by,

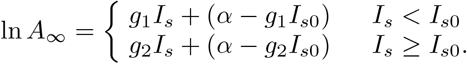

The f-I gain, *g*, was then defined to be the slope of the first linear line, *g* = *g*_1_, during the active condition. During the passive condition, the slope was approximately linear for all values of *I*_*s*_. As both the amplitude *D* and duration Δ_*D*_ of the dendritic spike increased, the two slopes merged, *g*_1_ ≈ *g*_2_, causing our definition of *g*_1_ to be undefined in the large amplitude and duration regime (Fig. 1g). The max coefficient of variation modulation, *c*, was defined as as the maximum difference in *C*_*v*_ between the active and passive conditions. For large amplitude *D* and duration Δ_*D*_, the peak *C*_*v*_ in the active condition moves to negative values of the somatic input current *I*_*s*_, which we did not include in our simulation. Therefore, we label this region as undefined, as we did not measure the peak in this regime.

### Experimental methods

Electrophysiological data was previously published [22]. Briefly, parasagittal slices of neocortex were prepared from young adult Wistar rats (P26-43). Recordings were performed at 34^*o*^C, targeting L5 pyramidal cells of the somtosensory cortex. Cells were chosen if their apical dendrites could be followed far enough to establish that the dendritic tree was intact. Patch electrode (5-10 MΩ for soma, 7-15 MΩ for dendrites) were made with an electrode puller (Sutter, USA, Model P-97). One electrode was always placed on the cell soma while simultaneously placing another electrode on the apical dendrite (445-719 *μ*m from the soma). Pipettes were filled with a solution containing (in mM): potassium gluconate 105, KCl 30, HEPES 10, MgCl_2_ 2, MgATP 2, Na_2_ATP 2, GTP 0.4 at pH 7.3. Naturalistic inputs were generated according to an Ornstein-Uhlenbeck process with correlation length of 3 ms and a noise amplitude of *σ* = 300 pA. The mean of the somatic current injection started at *I*_*s*_ = 0 pA and was increased in steps of Δ*I*_*s*_ = 50 pA which lasted for a duration of 2 s. With a total number of 20 steps. Experiments with this stepped somatic current injection were repeated with mean dendritic current injections of, *I*_*d*_ = (0, 250, 500, 750) pA.

### Data analysis

Experimental estimates of the single neuron firing rates and coefficient of variation were obtain by calculating the ISIs for each pair of somatic and dendritic mean input current. Each input level lasted 2s, allowing us to approximate the steady-state distribution. The firing rate of neuron *i* was then given by,

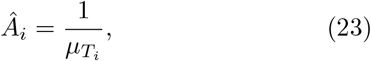

and the coefficient of variation of neuron *i* was given by,

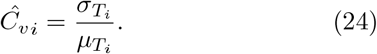

Where 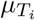 is the mean of the ISI distribution for for neuron *i* for a particular input pair, and 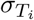 is the standard deviation. In Fig. 3b-d, we reported the average over *Â*_*i*_ and *Ĉ*_*v i*_, with errors bars representing the standard error in the mean.

## SUPPLEMENTAL

### Dendritic hotspots

The dendritic hazard rate given by equation (12) can be modified to account for spatial effects by adding a masking function *F* (*x*),

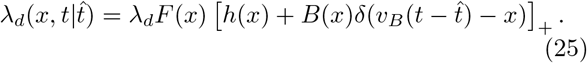

Where *F* (*x*) is a spatial function that captures the propensity for dendritic spikes to occur along the dendrite and is considered to be some intrinsic property of the dendrite, such as the density of ion channels. This can be used to bias dendritic spikes to the distal part of the dendrite (exponential filter) or concentrate dendritic spikes at a dendritic hotspot (Gaussian filter).

An example of a Gaussian hotspot with 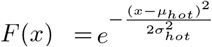 is shown in Fig. S1. The input currents have also been modified from Fig. 1 to be Gaussian; 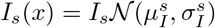 and 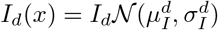. Note that the input potential in Fig. S1a is smoother than Fig.1 because of the Gaussian input. In Fig. S1a-c, the mean of the dendritic input is aligned with the mean of the dendritic hotspot, 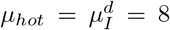. This gives rise to an effective input potential, 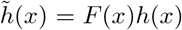, that is smaller than the original input potential. In Fig S1d-f, the dendritic input is unaligned with the hotspot, 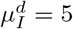. As a result, the modulation in gain and CV has been diminished.

Additional spatial properties of active dendrites can be explored by incorporating spatial dependence into how dendritic spikes effect the membrane potential. This can be accomplished by making the dendritic kernel in equation (7) explicitly depend on the location of the last den-dritic spike *x*^*′*^. For example, we could let the dendritic spike amplitude at the soma depend on the distance between the dendritic spike location and the soma. Lastly, local effects of dendritic spikes on the membrane potential, when *x >* 0, can also be added by modifying equation (7).

### Shunting inhibition

Using our formalism we can also investigate the case where the dendritic spike reaches the soma stochastically by adding a transmission probability distribution,

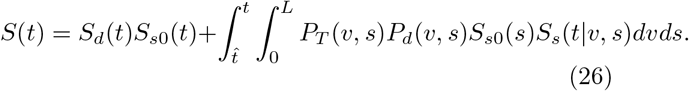

Where *P*_*T*_ (*x*^*′*^, *t*^*′*^) is the probability that the dendritic spike at point (*x*^*′*^, *t*^*′*^) successfully reaches the soma. Such a transmission probability distribution would capture any stochastic effect that inhibits dendritic spike propagation. An example would be random inhibitory inputs along the dendrite that shunt the dendritic spike.

---

**FIG. S1.**
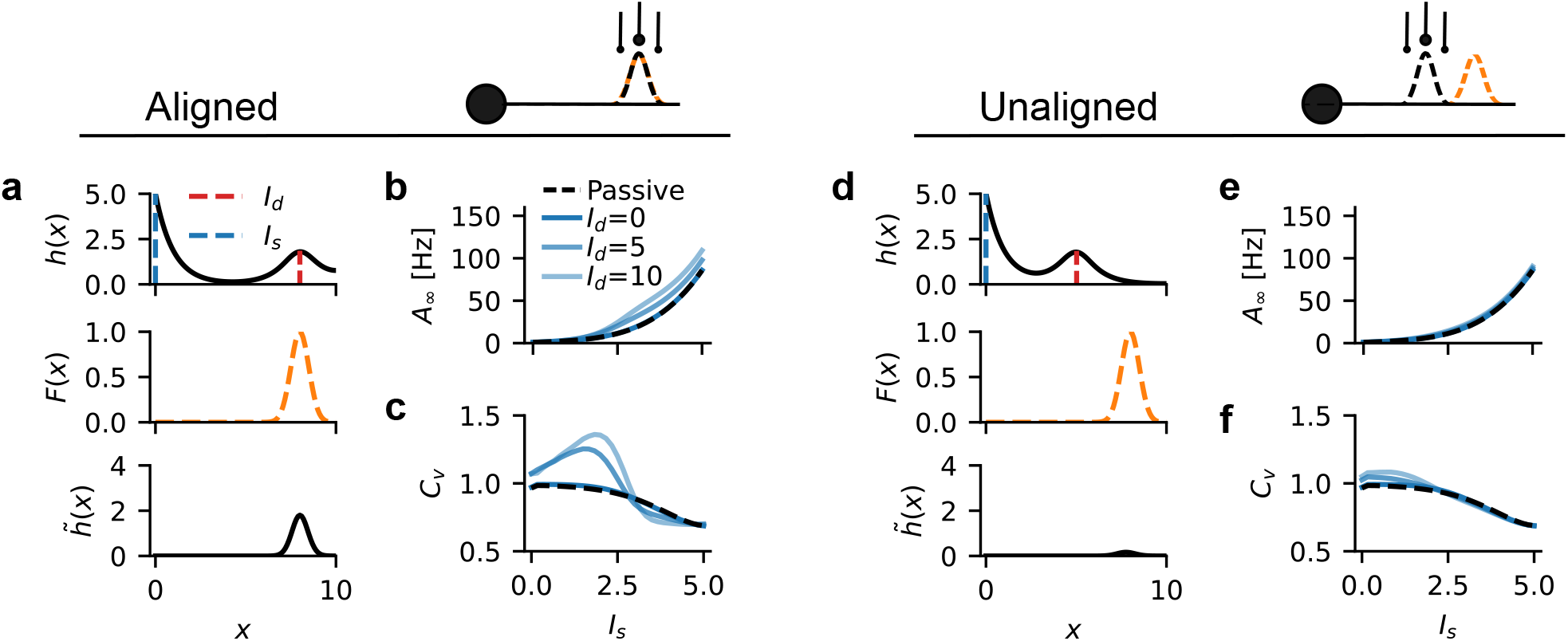
Generalization of the model to incorporate the spatial effects of a ‘hotspot’ for dendritic spike generation. **a-c** Dendritic ‘hotspot’ aligned with dendritic input. **a** Top: Stationary input potential resulting from the Green’s function of a ball-and-stick model receiving Gaussian distributed input currents at the soma 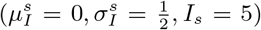 and in the dendrite 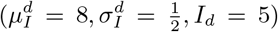. Middle: Gaussian dendritic hotspot 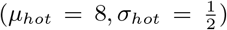. Bottom: Effective stationary input potential. **b** Firing rate and **c** coefficient of variation against somatic input current for three dendritic input strengths. The black dashed line corresponds to the absence of dendritic spikes. **d-f** Same as **a-c** but with the dendritic input unaligned with the dendritic ‘hotspot’ 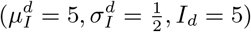.

## Notes

### Competing Interest Statement

The authors have declared no competing interest.

### Summary of Updates

The abstract, introduction, and discussion have been updated. Monte Carlo simulations have been added to Figure 1 and are explained in the methods section. A supplemental section has been added to the end of the manuscript.

